# Predicting future learning from baseline network architecture

**DOI:** 10.1101/056861

**Authors:** Marcelo G. Mattar, Nicholas F. Wymbs, Andrew S. Bock, Geoffrey K. Aguirre, Scott T. Grafton, Danielle S. Bassett

## Abstract

Human behavior and cognition result from a complex pattern of interactions between brain regions. The flexible reconfiguration of these patterns enables behavioral adaptation, such as the acquisition of a new motor skill. Yet, the degree to which these reconfigurations depend on the brain’s baseline sensorimotor integration is far from understood. Here, we asked whether spontaneous fluctuations in sensorimotor networks at baseline were predictive of individual differences in future learning. We analyzed functional MRI data from 19 participants prior to six weeks of training on a new motor skill. We found that visual-motor connectivity was inversely related to learning rate: sensorimotor autonomy at baseline corresponded to faster learning in the future. Using three additional scans, we found that visual-motor connectivity at baseline is a relatively stable individual trait. These results suggest that individual differences in motor skill learning can be predicted from sensorimotor autonomy at baseline prior to task execution.

**Highlights:**

- Sensorimotor autonomy at rest predicts faster motor learning in the future.
- Connection between calcarine and superior precentral sulci form strongest predictor.
- Sensorimotor autonomy is a relatively stable individual trait.

## 1 Introduction

Adaptive biological systems display a common architectural feature that facilitates evolvability (Kirschner and Gerhart, 1998; Kashtan and Alon, 2005; Félix and Wagner, 2008). That feature is modularity, or near-decomposability (Simon, 1965), in which the system is composed of small subsystems (or modules) that each perform near-unique functions. This compartmentalization reduces the constraints on any single module, enabling it to adapt to evolving external demands relatively independently (Kashtan and Alon, 2005; Wagner and Altenberg, 1996; Schlosser and Wagner, 2004). These principles relating modularity to adaptivity are evident across the animal kingdom, offering insights into phenomena as diverse as the developmental program of beak morphology in Darwin’s finches (Mallarino et al., 2011) and the heterochrony of the skeletal components of the mammalian skull (Koyabu et al., 2014).

While an intuitive concept in organismal evolution, where genetic programs drive dynamics over long time scales, it is less clear how modularity might confer functional adaptability in neural systems whose computations are inherently transient and fleeting. To gain conceptual clarity, we consider synchronization: a foundational neural computation that facilitates communication across distributed neural units (Fries, 2005; Voytek et al., 2015). Recent evidence from the field of statistical physics demonstrates that synchronization of a dynamical system is directly dependent on the heterogeneity of the associations between units (Gomez-Gardenes et al., 2007). Specifically, in systems where units with oscillatory dynamics are coupled in local modules, each module can synchronize separately (Arenas et al., 2006), offering the potential for unique functionality and independent adaptability. These theoretical observations become intuitive when we consider *graphs*: visual depictions of nodes representing oscillators, and edges representing coupling between oscillators (Fig. 1a). Modules that are densely interconnected will tend to become synchronized with one another, and each module will therefore be unable to adapt its dynamics separately from the other module (Arenas et al., 2006). This highly constrained state decreases the potential for adaptability to incoming stimuli in a changing environment. Conversely, modules that are sparsely interconnected with one another will maintain the potential for adaptive, near-independent dynamics.

**Figure 1:**
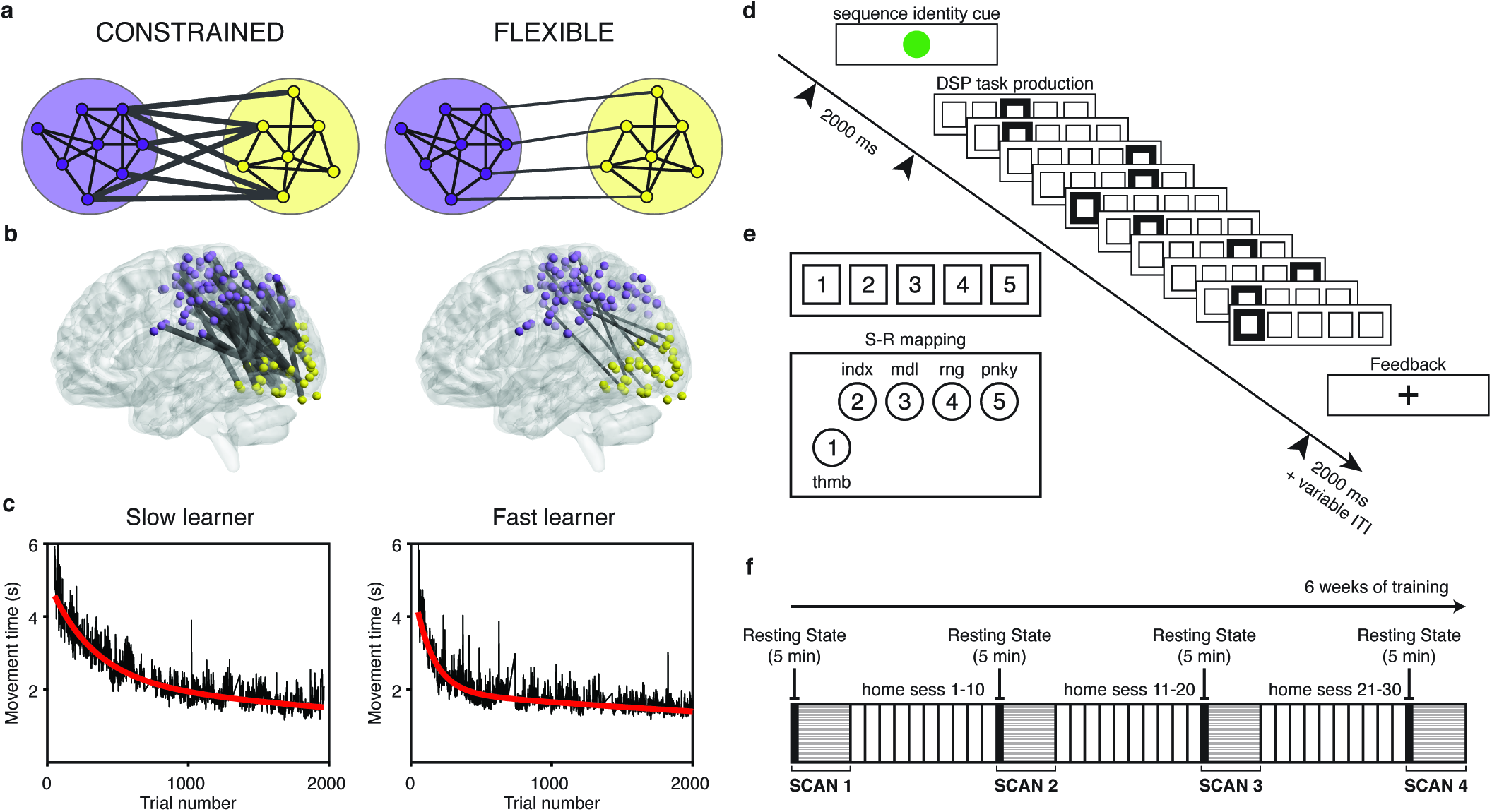
Network dynamics constrain adaptive learning behavior. (*a*) The degree of connectivity between two modules can impose important constraints on the types of dynamics that are possible. A lower degree of statistical dependence between the activity profiles of two modules can allow for greater flexibility in module dynamics. (*b*) Learning a new motor skill — a sequence of finger movements — induces a progressive change in the connectivity between visual and somato-motor cortices in humans (Bassett et al., 2015). We hypothesize that individuals who display a greater functional separation, or greater modularity, between motor and visual modules at rest are poised for enhanced adaptability, and therefore will learn faster over the 6 weeks of practice than individuals who display less functional separation between these modules. (*c*) Time in seconds required to correctly perform each sequence of finger movements (here referred to as *movement time*) for two example human subjects over 6 weeks of training. We observe an exponential decay in the trial-by-trial movement times for all participants (black lines), indicating that learning is occurring. The exponential drop-off parameter of a two-term exponential fit (red line) quantifies how rapidly each participant learned. Left and right panels illustrate the fits for an example slow and fast learner, respectively. (*d*) On each trial, the initial stimulus indicated which sequence should be performed. Each correct key press led to next stimulus cue until the ten-element sequence was correctly executed. At any point, if an incorrect key was hit, a participant would receive an error signal (not shown in the figure), and the sequence would pause until the correct response was received. (*e*) Stimulus-response mapping between a conventional keyboard or an MRI-compatible button box (lower left) and a participant’s right hand. (*f*) Training occurred over the course of 30 or more behavioral training sessions spanning approximately 42 days. Participants were scanned on the first day of the experiment and on three other occasions spaced approximately 1.5-2 weeks from one another. Each scan session began with a 5-minute resting state scan.

Given these theoretical observations in oscillator networks, we hypothesize that human brains display a modular architecture for the explicit purpose of facilitating behavioral adaptability (Meunier et al., 2010; Bullmore et al., 2009). Such a hypothesis is bolstered by evidence that neuronal cell distributions evolve differently in regions of the brain that code for simpler reflexive *versus* more complex adaptive functions (Lewitus et al., 2012). The hypothesis also has implications for individual differences in cognitive ability across humans. Specifically, we expect that individuals that display greater modularity, or sparser connectivity, between task-specific modules should also display more behavioral adaptability in the face of novel task demands (Bassett et al., 2011, 2013, 2015) (Fig. 1b). We expect that modularity should be particularly important between low-level modules that must evolve independently; connections involving higher-level control areas could have a different relationship due to the importance of these connections in the acquisition of new skills (Cole et al., 2013).

To test these hypotheses, we studied a cohort of healthy adult human subjects who learned a new motor skill from visual cues over the course of 6 weeks (Fig. 1c). During this timeframe, recorded fMRI activity during task execution shows that learning induces a growing autonomy between motor and visual systems (Bassett et al., 2015). Here, we focused on functional connectivity *at rest* acquired from the same cohort, prior to the onset of learning. We hypothesized that individuals who display a greater functional separation, or greater modularity, between motor and visual modules at rest are poised for enhanced adaptability in this task, and therefore should learn faster over the 6 weeks of practice than individuals who display less functional separation between these modules. Further, we ask whether this baseline *segregation* between modules is a *trait* of an individual, consistently expressed over multiple scanning sessions, or a *state* of an individual, and therefore potentially **responsive** to external manipulation or internal self-regulation. The answers to these questions have direct implications for predicting and manipulating a human’s ability to adapt its behavior — or learn — in the future.

The experimental protocol comprised of six weeks of training on six distinct motor sequences. Following a brief explanation of the task instructions, an initial MRI scan session was held during which blood-oxygen level dependent (BOLD) signals were acquired from each participant. The scan session began with a resting state scan lasting 5 minutes where participants were instructed to remain awake and with eyes open without fixation. During the remainder of the first scan session (baseline training), participants practiced each of six distinct motor sequences in a discrete sequence production (DSP) task for 50 trials each, or approximately 1.5 hours. They were then instructed to continue practicing the motor sequences at home using a training module that was installed by the experimenter (N.F.W.) on their personal laptops. Participants completed a minimum of 30 home training sessions, which were interleaved with two additional scan sessions, each occurring after at least 10 home training sessions. A final scan session was held following the completion of the 6 weeks of training. The same protocol was followed in each of the four scan sessions: a 5 minute resting state scan, followed by approximately 1.5 hours of the DSP task, where each of six distinct motor sequences was practiced for 50 trials each.

## 2 Results

### 2.1 Behavioral markers of learning

Participants practiced a set of ten-element motor sequences in a DSP paradigm (Fig. 1d). Training occurred over the course of 30 or more behavioral training sessions spanning approximately 42 days, for a total of over 2,000 trials (Fig. 1f; Fig. S1). The time required to correctly perform each sequence (movement time) decayed exponentially over time, and the rate of this decay displayed remarkable individual variability (Fig. 1c, Fig. S2). To quantify this feature of behavior, we defined the *learning rate* as the exponential drop-off parameter of the movement times, collated from home training sessions over the course of the entire experiment and averaged between two extensively practiced sequences (EXT sequences; see Methods) (Bassett et al., 2015). The learning rate — which quantifies how rapidly each participant converges to their own optimal performance — varied between 2.7 × 10^−3^ and 8.0 × 10^−3^ trial^−1^ (*M* = 5.2 × 10^−3^, *SD* = 1.6 × 10^−3^ trial^−1^). These data indicate that the fastest learner converged to relatively steady performance approximately three times faster than the slowest learner (Fig. 1c).

### 2.2 Sensorimotor initialization predicts future learning

Next, we asked whether a modular architecture during resting state - an important correlate of underlying structural connectivity (Honey et al., 2009; Goñi et al., 2014) and a marker of prior experience (Taylor et al., 2012; Duan et al., 2012; Burton et al., 2014) - is predictive of behavioral adaptability. To address this question, we considered a visual module and a motor module identified in a previous study with the same subjects (Bassett et al., 2015). These modules, derived from the analysis of fMRI data acquired during the performance of the motor sequence task, were shown to become less integrated with one another as sequence performance became more automatic (Bassett et al., 2015). We therefore hypothesized that functional connections between the same modules would, at baseline, explain individual variability in future learning rate.

To test this hypothesis, we analyzed the spontaneous fluctuations in BOLD activity during a 5-minute scan immediately prior to the initial task practice session. We parcellated the brain into a set of 333 functionally-defined regions representing putative cortical areas (Gordon et al., 2014) and identified the subset of these regions corresponding to the two modules of the task-based fMRI study with the same cohort (Bassett et al., 2015). These two sets of regions broadly corresponded to (i) early visual cortex (which has been referred to as the *visual module;* Fig. 2a; Table 1) and (ii) primary and secondary somatomotor regions (which has been referred to as the *somato-motor module;* Fig. 2a; Table 1).

**Figure 2:**
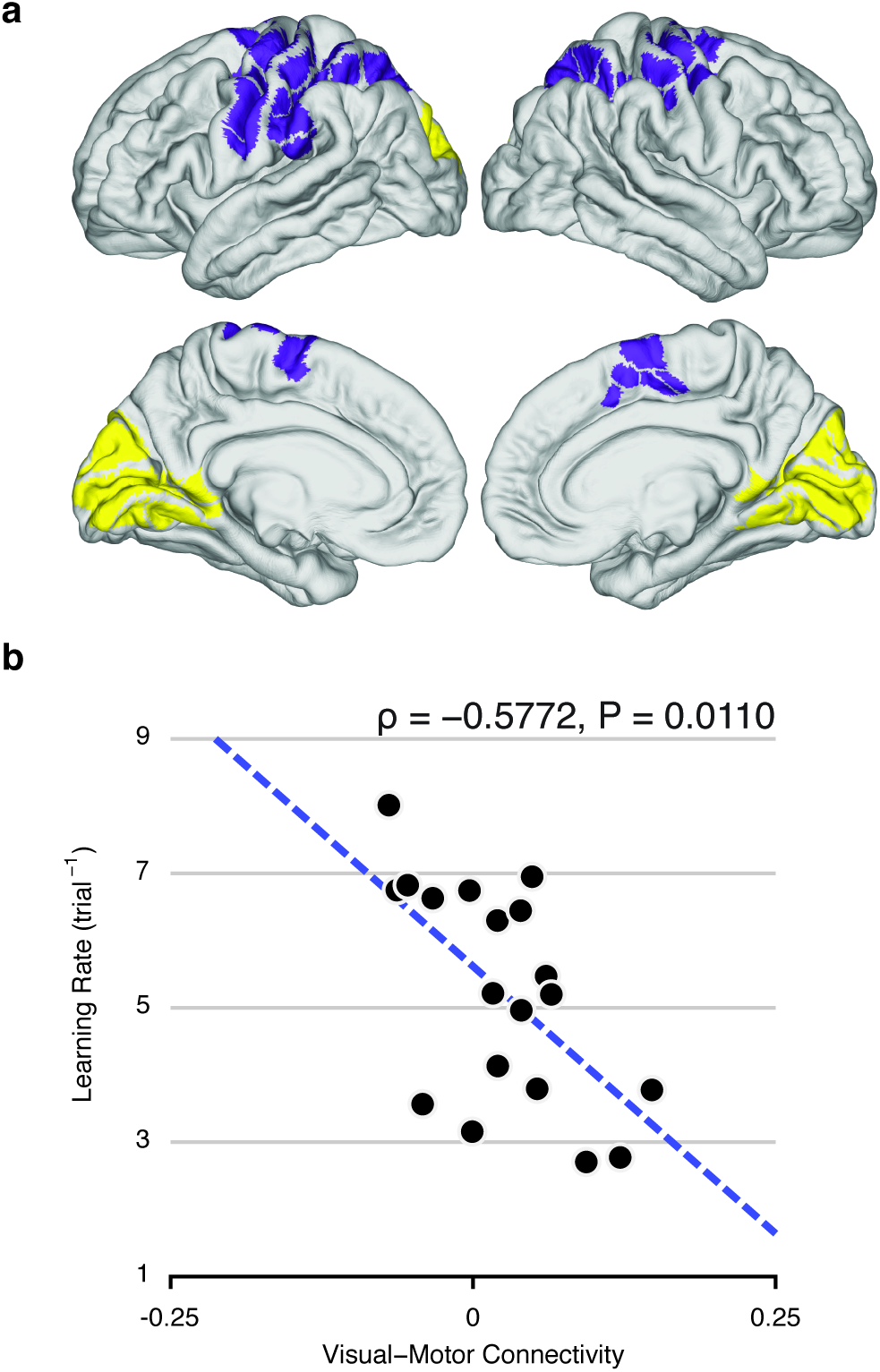
Baseline visual-motor connectivity predicts future learning rate. (*a*) Visual module (yellow) and somato-motor module (purple), identified by time-resolved clustering methods applied to BOLD activity acquired during execution of motor sequences (Bassett et al., 2015). The modules were defined in a data-driven manner and correspond broadly to putative visual and somato-motor modules. (*b*) Functional connectivity between visual and somato-motor modules, estimated at rest and prior to learning, reliably predicts individual differences in future learning rate. We define the learning rate as the exponential drop-off parameter of the participant’s movement time as a function of trials practiced, and we define functional connectivity as the average correlation value between activity in visual regions and somato-motor regions. Note that we use the term “prediction” to imply that the value of one variable (at one point in time) can be used to predict the value of another variable (at a later point in time), without implying the use of out-of-sample generalization (Gabrieli et al., 2015).

**Table 1:**
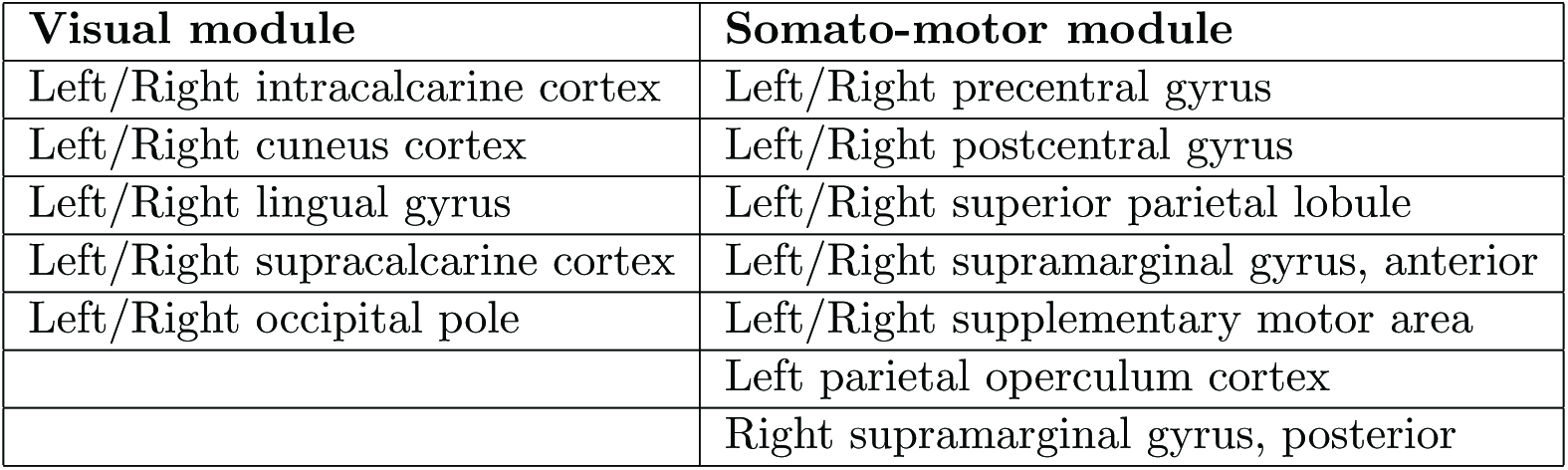
Brain areas in visual and somato-motor modules.

We then asked whether the interactions between these two modules at baseline were predictive of future learning rate. We extracted the average resting state time series from each brain region and calculated their pairwise Pearson correlation coefficient. Next, we applied a Fisher *z*-transform to these coefficients and calculated the average *z*-transformed correlation between regions in the visual and somato-motor modules. We refer to this value as the *visual-motor connectivity.* We observed that individuals with low visual-motor connectivity at rest, prior to any task practice, exhibited a larger learning rate in the following 6 weeks of practice (Spearman’s rank correlation: *ρ* = −0.5772, *P* = 0.0110; Fig. 2b). Similar results were obtained using an anatomically-defined parcellation with 626 regions (Spearman’s rank correlation: *ρ* = −0.6211, *P* = 0.0055; Fig. S3). These results suggest that baseline visual-motor connectivity can be thought of as a sensorimotor initialization parameter that constrains adaptive learning behavior.

To confirm that these task-based modules were also effective modules at rest, we calculated the modularity quality of this partition during the resting state (Equation (2) in Methods). The value obtained for this partition (*Q* = 0.4226 ± 0.0719 SEM) was larger than the modularity of all 10,000 random partitions of visual and motor regions (*Q* = −0.0108, CI: [-0.0310,0.0256], *P* = 0.0001) and also larger than the modularity of all 10,000 random sets of the modules in the brain of equal size to visual and motor regions (*Q* = −0.0106, CI: [-0.0313,0.0270], *P* = 0.0001). Therefore, the visual and somato-motor modules used in our analyses are also effective modules at rest.

### 2.3 Behavioral and neural specificity

The relationship between resting visual-motor connectivity and future behavior was highly specific to learning rate, being unrelated to error rates, reaction time, or other parameters of the fitted movement time *versus* trials-practiced curve (Fig. S4). Moreover, the relationship remained significant (*α* = 0.05) even after regressing out the effect of initial performance (Spearman’s rank correlation: *ρ* = −0.5614, *P* = 0.0138) or after regressing out the effects of both initial and final performances (Spearman’s rank correlation: *ρ* = −0.4684, *P* = 0.0448), and became marginally significant after regressing out the effect of final performance (Spearman’s rank correlation: *ρ* = −0.4526, *P* = 0.0533). Therefore, baseline visual-motor connectivity is specifically related to the rate of decay of movement time (*learning rate*).

Having established that baseline functional connectivity between broadly defined visual and somatomotor areas predicts individual differences in future learning rate, we next explored which specific subregions — or functional connections — within visual and somato-motor areas might be most responsible for driving this effect. For this analysis, we used an annotated surface-based parcellation (`aparc.a2009s.annot` in Freesurfer) which has a label for each cortical region (Destrieux et al., 2010). We observed a general trend for negative correlations between visual-motor connectivity and learning rate, as evident from the predominantly blue color in Fig. 3a (Spearman’s rank correlation between visual-motor connectivity and learning rate, using broad visual and somato-motor regions of interest from a surface-based parcellation, was: *ρ* = −0.5596, *P* = 0.0141). This result indicates that the broader regions selected in surface space still retain the overall properties of the original parcellation with task-identified modules. To test whether some functional connections were significantly more correlated with learning rate than others, we used a bootstrap procedure with 10,000 subject samples with replacement to derive the sampling distribution of each correlation value in Fig. 3a. We observed that individual differences in future learning rate were most strongly predicted by functional connectivity between the premotor area adjacent to the right superior precentral sulcus and early-visual areas adjacent to the calcarine sulcus in both hemispheres (Left calcarine sulcus to right superior precentral sulcus: Spearman’s *ρ* = −0.8211, bootstrap: *M* = −0.7935, 95% *CI* = [−0.9365,−0.5434]; Right calcarine sulcus to right superior precentral sulcus: Spearman’s *ρ* = −0.8228, bootstrap: *M* = −0.7904, 95% *CI* = [−0.9043,−0.6060]; Fig. 3b). Across all bootstrap samples, these two values were larger than 98% of the others, demonstrating that these connections are robustly more correlated with learning rate than other visual-motor connections.

**Figure 3:**
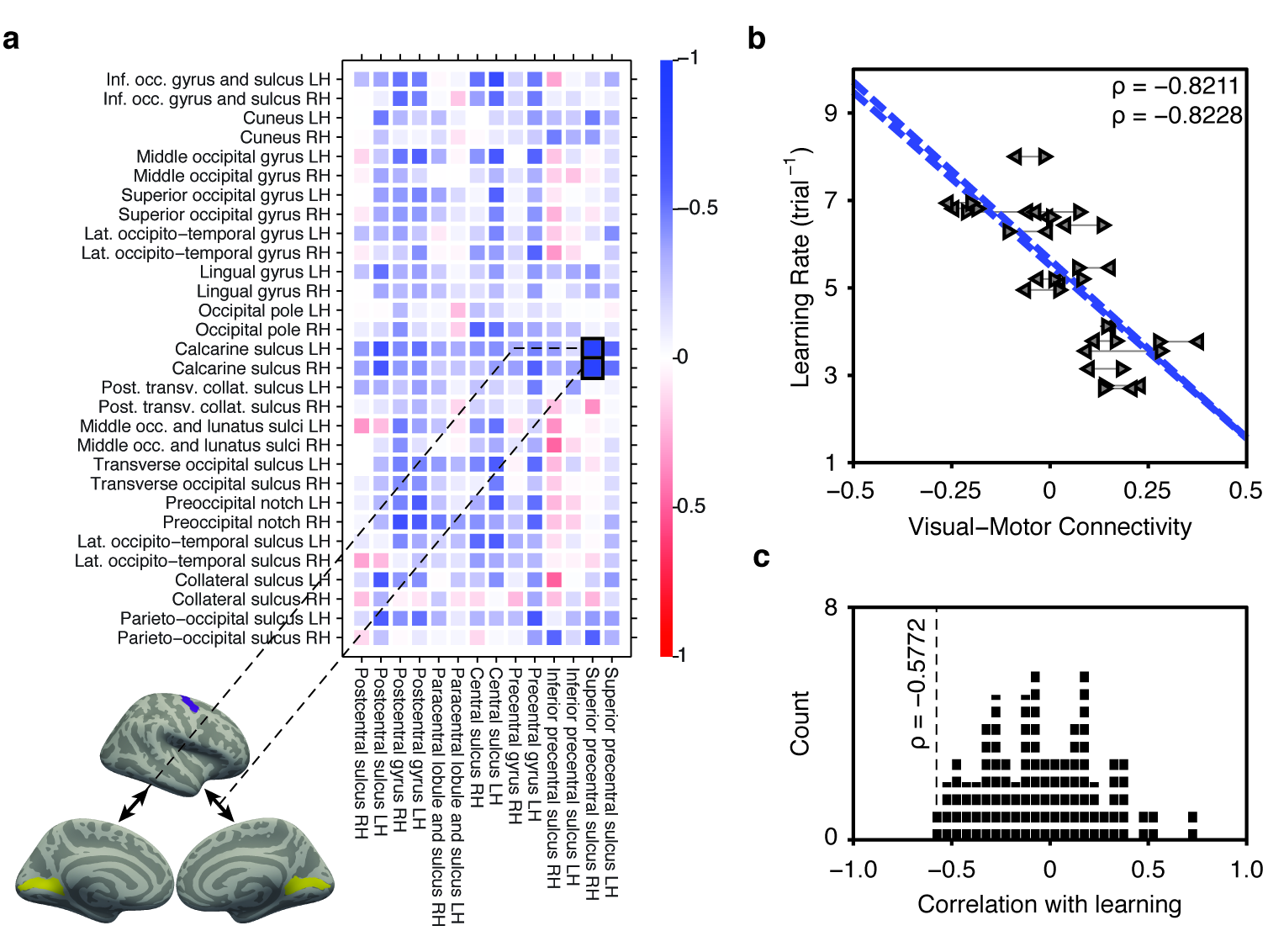
Learning rate is best predicted by connectivity between early visual and dorsal premotor areas. (*a*) Using a surface-based annotation encompassing broadly defined visual and somatomotor areas, we calculated the correlation between learning rate and the functional connectivity between each pair of subregions (negative correlations are represented in blue; positive correlations are represented in red). Learning rate was best predicted by connectivity between early-visual areas adjacent to the calcarine sulcus in both hemispheres (yellow) and the dorsal premotor area adjacent to the right superior precentral sulcus (purple). (*b*) Functional connectivity between left calcarine sulcus and right superior precentral sulcus significantly predicted individual differences in future learning rate (data points are indicated by left pointing triangles). Similarly functional connectivity between right calcarine sulcus and right superior precentral sulcus significantly predicted learning rate (data points are indicated by right pointing triangles). (*c*) Distribution of correlation values between learning rate and module-to-module connectivity across subjects. Visual-motor connectivity has one of the highest correlations with learning rate.

We then wished to examine whether the observed correlation between visual-motor connectivity was specific to visual and motor modules, or whether this effect was also present in other regions of the brain. We considered a set of 12 putative functional modules assigned to various regions of the parcellation used: auditory, cingulo-opercular, cingulo-parietal, default mode, dorsal attention, fronto-parietal, retrosplenial-temporal, somato-motor hand, somato-motor mouth, salience, ventral-attention, and visual (Gordon et al., 2014). We then calculated the average pairwise connectivity between each pair of putative modules, and the correlation between learning rate and module-to-module connectivity across subjects. Using this approach, we observed that connectivity between our task-derived modules was one of the most predictive of learning rate (2nd out of 66 pairs, *P* = 0.0299; Fig. 3c). This suggests that the relationship between modularity and learning rate was highly specific to visual-motor connectivity.

Nonetheless, prior evidence suggests a critical role for online cognitive control during learning (Galea et al., 2010; Chrysikou et al., 2014) and adaptive behavior in general (Chrysikou et al., 2011; Thompson-Schill et al., 2009). Recent analyses using graph theory suggest that at least five distinct modules are associated with cognitive control (Power et al., 2011)): the fronto-parietal network, the cingulo-opercular control network, the salience network, the ventral attention network and the dorsal attention network. We therefore examined the degree to which connectivity within these five modules correlated with learning rate. We found that connectivity within the cingulo-opercular network was the only one that significantly correlated with learning rate (Spearman’s rank correlation: *ρ* = −0.6228, *P* = 0.0053, Bonferroni adjusted *p*-value: *P* = 0.0265), although the exploratory character of these analyses suggests that they be interpreted as preliminary evidence.

Finally, we wished to verify whether visual-motor connectivity at baseline is related to other network-derived metrics. In particular, network flexibility has been previously shown to predict the learning rate in future sessions in a similar motor learning paradigm using cued rather than discrete sequence production (Bassett et al., 2011). Flexibility is defined as the proportion of times in which a given node changes module affiliation (Bassett et al., 2011). Using functional connectivity data acquired during task execution, we calculated the average node-wise flexibility for each subject and each session. We then computed the correlation between visual-motor connectivity and flexibility. We observed that visual-motor connectivity was uncorrelated with network flexibility on the first scan (Spearman’s rank correlation: *ρ* = 0.0240, *P* = 0.9224), as well as with the avearge network flexibility across scans (Spearman’s rank correlation: *ρ* = 0.0714, *P* = 0.7716). We also assessed the degree to which visual-motor connectivity predicts learning rate controlling for network flexibility. We found that visual-motor connectivity still predicted learning rate when controlling for network flexibility on the first scan (partial Spearman’s rank correlation: *ρ* = 0.6020, *P* = 0.0082) and when controlling for avearge network flexibility across scans (partial Spearman’s rank correlation: *ρ* = 0.5846, *P* = 0.0108). Therefore, visual-motor connectivity predicts learning rate independently from network flexibility.

### 2.4 Sensorimotor initialization: A state or a trait?

Given the predictive nature of baseline visual-motor connectivity, one might wish to know whether this baseline varies from day to day, thereby playing the role of an online initialization system, or whether it remains relatively stable over the course of the 6-week experiment. That is, are we measuring a network property related to learning that varies from session to session (over the course of hours or days) or is this a consistent relationship over the entire experiment, indicative of a trait effect? The answer to this question could offer much needed insight into the potential neurophysiological mechanisms underlying the observed relationship between baseline connectivity and learning: for example, from stable trait markers of structure (Honey et al., 2009; Goni et al., 2014) or prior experience (Taylor et al., 2012; Duan et al., 2012; Burton et al., 2014) to dynamic state markers of arousal (Nassar et al., 2012).

To address this question, we examined data from the three additional resting state sessions obtained throughout the 6 week training period (Fig. 1f). Therefore, a total of 4 resting state scan sessions separated by 1.5-2 weeks, each lasting 5 minutes, were examined for each subject. We then conducted a repeated measures ANOVA across the four scans and examined the sources of variance. We observed no consistent trend in the evolution of visual-motor connectivity across sessions, with only 5.5% of the total variance being explained by session (*F*(3, 54) = 1.7710, *P* = 0.1637). In contrast, 38.2% of the total observed variance in visual-motor connectivity was accounted for by differences between subjects (*F*(18, 54) = 2.0352, *P* = 0.0231). These observations suggest the existence of a significant *trait* marker.

How does the trait *versus* state nature of visual-motor connectivity impact prediction accuracy? When estimating the stable trait component by averaging an individual’s visual-motor connectivity values over all four scanning sessions, we observed that this trait component significantly predicts learning rate over the 6 weeks of training (Spearman’s *ρ* = −0.4614, *P* = 0.0484; Fig. 4b). When using the median visual-motor connectivity as an estimator for the trait component, however, the relationship was no longer significant (Spearman’s *ρ* = −0.3579, *P* = 0.1329). Importantly, there is clearly additional variance that is not explained by this trait component, as evidenced by session-to-session variability in visual-motor connectivity (Fig. 4a). Indeed, the relationship between visual-motor connectivity in the first session and learning rate remained significant even after regressing out the average trait component from each individual’s visual-motor connectivity (Spearman’s rank correlation: *ρ* = −0.4772, *P* = 0.0405). Thus, we hypothesized that session-to-session fluctuations in visual-motor connectivity could also explain the amount of learning on a session-to-session basis.

**Figure 4:**
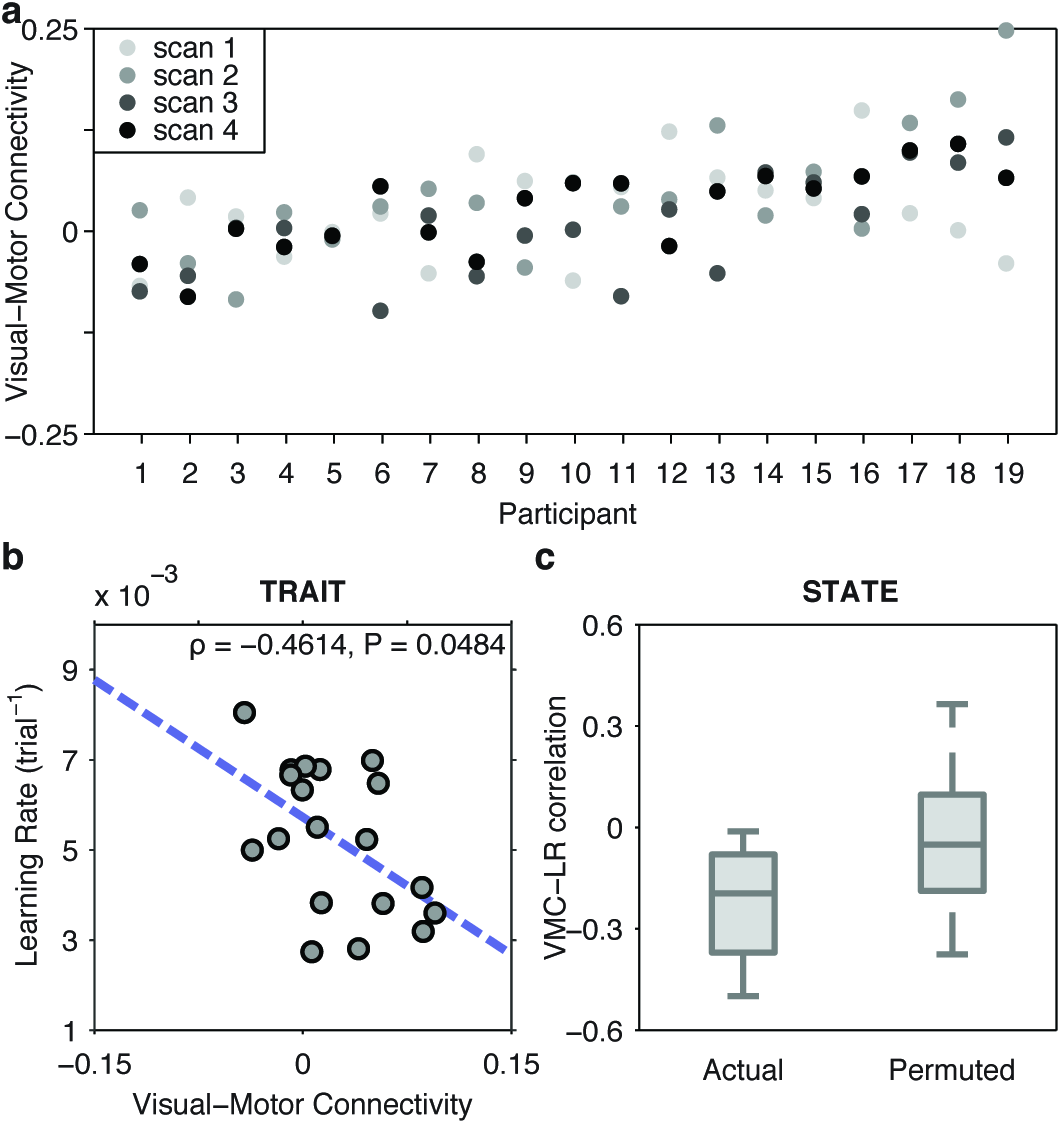
Visual-motor connectivity as a trait and as a state. (*a*) Between-session variability of visual-motor connectivity. For each participant, dots represent visual-motor connectivity measured at each of four resting state scans conducted immediately prior to task execution. Despite large variability between sessions, 38.2% of the observed visual-motor connectivity variance was accounted for by a trait marker representing between-subject variability. (*b*) The *trait* marker is the component of visual-motor connectivity that remains stable across time, with the variability from session to session here termed the *state* component. The average visual-motor connectivity across all four sessions, an estimator of the trait component of visual-motor connectivity, significantly predicted overall learning rate (*ρ* = −0.4614, *P* = 0.0484). (*c) Left:* Spearman’s correlation coefficients between session-specific learning rate, estimated from trials performed inside the scanner immediately following resting state scans, and session-specific visual-motor connectivity. *Right:* Spearman’s correlation coefficients for all 24 permutations of resting state scans to task sessions, between visual-motor connectivity and session-specific learning rates. The actual pairing of resting state scans to task sessions had the strongest average correlation from all possible pairings (*P* = 0.0400), indicating that the state component of visual-motor connectivity has some degree of temporal specificity.

To assess the potential predictive role of state dependent components of visual-motor connectivity, we asked whether visual-motor connectivity estimated from a single baseline scan predicts learning rate in a temporally adjacent training session more so than in temporally distant training sessions. To estimate a session-specific learning rate, we used movement times from minimally trained sequences (MIN) to ensure learning (as indexed by a reduction in movement times) was still occurring throughout all four sessions. These trials were performed during scan sessions, in runs immediately following the resting state scans. While the individual correlations between session-specific learning rate and session-specific visual-motor connectivity were not statistically significant, their average (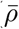 = −0.2261) was the largest of all possible pairings of resting state scans and task execution sessions (24 permutations, *P* = 0.0400). These results suggest that visual-motor connectivity contains both a trait and a state component, the former predicting a stable task aptitude and the latter predicting temporally-specific measures of learning.

## 3 Discussion

The understanding of many higher-level cognitive functions often requires one to study the brain during effortful thought (Gazzaniga and Mangun, 2014). Yet, some basic organizational principles and constraints can also be observed while the brain idles at baseline. Consistent evidence from multiple imaging modalities and subject cohorts demonstrate that the brain’s resting baseline is characterized by a modular (Meunier et al., 2009; Bullmore et al., 2009; Sporns and Betzel, 2015), or near-decomposable nature (Simon, 1965), and that these modules are composed of brain regions that tend to perform similar cognitive functions (Salvador et al., 2005; Power et al., 2011; Yeo et al., 2011; Cole et al., 2014). Yet, how this modular architecture supports the sequential and dynamic integration of the many high-level cognitive functions required during motor skill learning remains far from understood (Medaglia et al., 2015). Here we observe that individuals who display lower values of correlation between their resting baseline activity in motor and visual regions learn faster in the following 6 weeks of task practice. That is: our results suggest that a more modular architecture in low-level visual and motor regions may be beneficial for learning a visual-motor task. This result complements both empirical and theoretical lines of inquiry recently demonstrating that modular architecture confers robustness as well as evolvability simultaneously (Anderson and Finlay, 2014), helps organisms evolve new skills without forgetting old skills (Ellefsen et al., 2015), and – in the motor-visual system – increases as learning occurs (Bassett et al., 2015).

### 3.1 The Benefits of Independence

While the baseline separation between the entire motor and visual modules was predictive of individual differences in future learning behavior over 6 weeks of task practice, we also observed that the regional associations that drove this prediction most were the functional connections between the contralateral superior precentral sulcus and the bilateral calcarine sulcus. In classical models of motor processing and control, the superior precentral sulcus is thought of as the dorsal premotor area (Hardwick et al., 2013), and activation in this area is related to the performance of visual-motor hand/arm conditional responses (Amiez et al., 2006). It is well know that this region plays a central role in mapping visual cues to spatial motor responses in both human and non-human primates (Astafiev et al., 2003; Rushworth et al., 2003; Grefkes and Fink, 2005; Halsband and Lange, 2006; Kravitz et al., 2011). Given this specific role in motor-visual integration, it is interesting that individuals with the weakest baseline connections between this area and early visual cortices learn the fastest. One simple interpretation of these findings builds on the notion that the learning process is one in which the task of the brain is to develop direct motor-motor associations (Verwey, 2001; Wymbs et al., 2012): each finger movement directly triggers the next, without the need for visual cues. Individuals with low connectivity between dorsal premotor and visual areas – and therefore more independence or autonomy of visual and motor processes (Bassett et al., 2015) – are able to develop motor-motor associations faster.

Such an explanation suggests the presence of a broader competitive process that may play a role in other cognitive tasks: individuals that display greater integration between cognitive processes at rest may be less able to disengage such processes from one another during task execution. This hypothesis is indeed supported by preliminary evidence in both healthy and clinical cohorts. For example, in healthy adult subjects, increased modularity (decreased integration) of resting state functional connectivity networks has been shown to be positively correlated with improvement in attention and executive function after cognitive training (Arnemann et al., 2015). Similarly, individuals with greater negative correlation between default mode and working memory networks exhibited better behavioural performance on a working memory task (Sala-Llonch et al., 2012). Conversely, in subcortical vascular mild cognitive impairment, increased integration between modules in the inferior and superior parietal gyrus at rest has been shown to be associated with impaired cognitive performance (Yi et al., 2015). Finally, such a broad competitive process is supported by recent work in normative neurodevelopment showing that individuals with weaker sensorimotor integration at rest tended to display better cognitive performance (*N* = 780 in the Philadelphia Neurodevelopmental Cohort) (Gu et al., 2015).

### 3.2 Drivers of Baseline Architecture

A growing literature demonstrates the absolutely fundamental role of baseline network architecture in explaining individual differences in cognition and behavior. The strength of individual functional connections, or larger sets of connections, has been observed to correlate with individual differences in IQ (Song et al., 2008, 2009), fluid intelligence (Smith et al., 2013; Finn et al., 2015), attention (Rosenberg et al., 2016; Kessler et al., 2016; Poole et al., 2016), visual orientation discrimination (Baldassarre et al., 2012), working memory (Sala-Llonch et al., 2012; Zou et al., 2013), color knowledge (Wang et al., 2013), auditory stimulus detection (Sadaghiani et al., 2015), pursuit rotor performance (Wu et al., 2014), and the ability to learn foreign sounds (Ventura-Campos et al., 2013) and probabilistic regularities (Stillman et al., 2013). Yet, it is unclear what neurophysiological or develpmental factors drive these individual differences at baseline.

Current theories of resting state drivers can be summarized along two key dimensions: genetically-encoded structure, and prior or current experience. First, resting state functional connectivity is related to some degree to underlying large-scale structural connectivity as estimated by white matter tractography (Honey et al., 2009; Hermundstad et al., 2013, 2014; Goñi et al., 2014; Shen et al., 2015): two brain areas that are connected by a large number of white matter streamlines also tend to display strong correlations in their resting BOLD activity. These structural patterns may form a constraint on resting state dynamics, at least partially driven by the genetic codes underlying module formation (Richiardi et al., 2015). Yet, structural connectivity can only be a partial explanation, as resting state functional connectivity varies appreciably over time scales in which structure remains constant (Andellini et al., 2015; Deuker et al., 2009; Hutchison et al., 2013; Leonardi et al., 2014). It will be interesting in future to determine whether structural differences among individuals might explain some of the predictive relationship between resting state functional connectivity and future learning behavior.

The second key driver of resting state functional connectivity is experience. Over short time scales, resting state patterns are altered for up to 20 minutes following task performance (Barnes et al., 2009), being modulated by cognitive processes as diverse as short term memory (Gerraty et al., 2014) and visual-motor learning (Albert et al., 2009). Moreover, resting state connectivity can be altered over longer time scales with cognitive training (Arnemann et al., 2015), mindfulness training (Taylor et al., 2012; Taren et al., 2015), progressive neurological disorders (Pievani et al., 2011), and aging (Betzel et al., 2014). While recent and more distant experience can play a role, perhaps the more tantalizing observation is that a person’s arousal state is also directly linked to their resting state functional connectivity (Eilam-Stock et al., 2014). This finding is particularly interesting in light of our results from the state-trait analysis, which suggest that visual-motor connectivity is more correlated with learning occurring in the immediately following trials than with trials performed in a different session. The existence of these state-dependent predictors of future learning is consistent with recent observations that arousal systems may directly regulate learning by coordinating activity in the locus coeruleus and anterior cingulate cortex (Nassar et al., 2012). Future work is necessary to determine the degree to which arousal state – as opposed to prior training – might manipulate the pattern of resting state connectivity, priming the system to optimally learn in the immediate future.

### 3.3 Baseline Initializations vs. Transient, Online Control

Cognitive control is a critical driver of learning during task performance (Dumontheil, 2014; Galea et al., 2010; Dixon and Christoff, 2014; Bassett et al., 2015). In an exploratory analysis in our data, we observed that baseline functional connections within the cingulo-opercular network significantly correlate with individual differences in future learning. This finding provides preliminary evidence of the relative importance of (i) baseline architecture, which represents the initialization of the brain, and (ii) task-elicited dynamics, which represents transient, online control. In combination with prior literature, our results suggest that the relative autonomy of sensorimotor systems and the recruitment of the cingulo-opercular network at rest strengthens the motor-motor associations that enable automatic performance (Verwey, 2001; Wymbs et al., 2012).

### 3.4 Methodological Considerations

There are several important methodological and conceptual considerations relevant to this work. First, while we use the term *modularity*, we do not mean the traditional notion of pure encapsulation of function as propounded by Fodor in his historic contribution to the field: “Modularity of Mind” (Fodor, 1983). Instead, we use the term as mathematically defined in (Newman, 2006b) to mean separation or segregation without requiring complete independence. Second, it is important to be clear about what the estimate of learning rate used here measures and what it does not measure. Critically, the learning rate is independent of initial performance, a measurement of experience on similar tasks, and is independent of final performance, a measurement of finger mechanics. Finally, in this work, we utilize large-scale non-invasive human recording of BOLD signals. It would be interesting in future to determine whether the sensorimotor autonomy that we describe here is related to competitive sensorimotor interactions reported at the neuronal level (Grent et al., 2014).

### 3.5 Implications for Educational and Clinical Neuroscience

We have shown that baseline visual-motor connectivity is a strong predictor of learning rate specifically in a DSP paradigm, but it is possible that these results would generalize to other motor skills, or that baseline separation between relevant cognitive systems is, in general, beneficial for other classes of learning in perceptual, cognitive, or semantic domains. Predicting individual differences in future learning has massive implications for neurorehabilitation (in those who are aging, injured, or diseased) and neuroeducation (in children or older trainees). Predictors drawn from behavioral performance or from brain images acquired during behavioral performance necessarily have limited applicability in rehabilitation and education domains where subjects may be unable to perform the task, or be unable to lie still in a scanner during task performance. Predictors drawn from resting state scans offer the possibility for direct translation to the clinic and classroom. Moreover, our delineation of state and trait components of sensorimotor initialization predictors suggests the possibility of directly manipulating subject state, for example with non-invasive stimulation (Galea et al., 2010; Luber and Lisanby, 2014), neurofeedback (Bassett and Khambhati, 2017), or task priming (Enriquez-Geppert et al., 2013) to enhance future performance, thereby optimizing rehabilitation or training.

## 4 Materials and Methods

### 4.1 Participants

Twenty-two right-handed participants (13 females and 9 males; mean age of 24 years) volunteered to participate in this study. All volunteers gave informed consent in writing, according to the guidelines of the Institutional Review Board of the University of California, Santa Barbara. Three participants were excluded: one failed to complete the experiment, one had excessive head motion, and one had a functional connectivity profile whose dissimilarity to those obtained from other participants was more than three standard deviations away from the mean, potentially due to sleep (Fig. S6). The final cohort included 19 participants who all had normal or corrected vision and no history of neurological disease or any psychiatric disorder.

### 4.2 Experimental setup and procedure

In a discrete sequence-production (DSP) task, participants practiced a set of ten-element motor sequences, responding to sequential visual stimuli using their right hand (Fig. 1d). The visual display contained a horizontal array of five square stimuli, each corresponding to one finger. Mapped from left to right, the thumb corresponded to the leftmost stimulus and the smallest finger corresponded to the rightmost stimulus. The square corresponding to the current button press was highlighted in red, changing to the next square immediately following a correct button press. Only correct button presses advanced the sequence, and the time for completion was not limited. Participants were instructed to respond quickly and to maintain accuracy.

Six different ten-element sequences were used in the training protocol, with three possible levels of exposure: two sequences were extensively trained (EXT; 64 trials per session); two sequences were moderately trained (MOD; 10 trials per session); and two sequences were minimally trained (MIN; 1 trial per session). The same sequences were practiced by all participants. In each sequence, each of the five possible stimulus locations was presented twice and included neither immediate repetitions (e.g. “1-1”) nor regularities such as trills (e.g., “1-2-1”) or runs (e.g., “1-2-3”). A sequence-identity cue indicated, on each trial, what sequence the participant was meant to produce: EXT sequences were preceded by either a cyan (EXT-1) or a magenta (EXT-2) circle, MOD sequences were preceded by either a red (MOD-1) or a green (MOD-2) triangle, and MIN sequences were preceded by either an orange (MIN-1) or a white (MIN-2) star. No participant reported any difficulty viewing the identity cues. The number of error-free sequences produced and the mean time required to complete an error-free sequence was presented after every block of ten trials. See Fig. S7 for the number of trials performed for each sequence type.

Participants were scanned on the first day of the experiment (scan 1) and on three other occasions (scans 2–4) spaced approximately 1.5–2 weeks apart from one another. The entire experiment spanned approximately 42 days (Fig. S1). A minimum of ten home training sessions was completed in between any two successive scanning sessions, for a total of at least 30 home sessions. Home training sessions were performed on personal laptop computers using a training module installed by the experimenter.

Before the first scanning session, the experimenter provided a brief introduction to participants in which he explained the mapping between the fingers and the DSP stimuli, as well as the significance of the identity cues. Next, fMRI data was acquired as subjects rested quietly in the scanner prior to any task performance. Finally, fMRI data was acquired as subjects performed a series of trials on the DSP task spread over five scan runs, using a 5-button response box with distances between keys similar to placement on a standard 15 in laptop. Each scan run acquired during task performance contained 60 trials grouped in blocks of ten, and similarly to home training sessions, performance feedback was given at the end of every block. Each block contained trials belonging to a single exposure type (EXT, MOD or MIN), and included five trials for each of the two sequences. Therefore, an equal number of trials from each sequence was performed during scan sessions (50 trials per sequence, for a total of 300 trials per scan session; Fig. S7). Trial completion was indicated by a fixation cross, which remained on the screen until the onset of the next sequence identity cue (the intertrial interval varied between 0 s and 6 s).

Two sessions were abbreviated due to technical challenges. In each case when a scan was cut short, participants completed four out of the five scan runs for a given session. We included behavioral data from these abbreviated sessions in this study.

### 4.3 Behavioral apparatus

In home train sessions, stimuli were presented with Octave 3.2.4 and Psychtoolbox 3 (Brainard, 1997) on each participants’ laptop computer. During scan sessions, stimuli were presented with MATLAB version 7.1 (Mathworks, Natick, MA) and Psychtoolbox 3 (Brainard, 1997), backprojected onto a screen and viewed through a mirror. Key presses and response times were collected using a custom fiber optic button box and transducer connected via a serial port (button box, HHSC-1 × 4-l; transducer, fORP932; Current Designs, Philadelphia, PA), with design similar to those found on typical laptops. For instance, the center-to-center spacing between the buttons on the top row was 20 mm (compared to 20 mm from “G” to “H” on a recent version of the MacBook Pro), and the spacing between the top row and lower left “thumb” button was 32 mm (compared to 37 mm from “G” to the spacebar on a MacBook Pro).

### 4.4 Behavioral estimates of learning

Following standard conventions in this literature, we defined the movement time (*MT*) as the difference between the time of the first button press and the time of the last button press in a single sequence. We calculated *MT* for every sequence performed in home training sessions over the course of the 6 weeks of practice. Across all trials in home training sessions, the median movement time was, on average, 1.70 s (average minimum 1.03 s and average maximum 7.12 s), with an average standard deviation of 0.79 s. For each participant and each sequence, the movement times were fit with a two-term exponential model (Schmidt and Lee, 1988; Rosenbaum, 2009) using robust outlier correction (operationalized by MATLAB’s function “fit.m” in the Curve Fitting Toolbox with option “Robust” and type “LAR”), according to Equation (1):

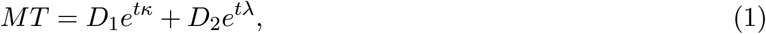

where *t* is time, *κ* is the exponential drop-off parameter (which we refer to as the *learning rate*) used to describe the fast rate of improvement, λ is the exponential drop-off parameter used to describe the slow, sustained rate of improvement, and *D*_1_ and *D*_2_ are real and positive constants. The magnitude of *κ* indicates the steepness of the learning curve: curves with larger *κ* values decay more quickly than curves with smaller *κ* values. Therefore, *κ* indicates the speed of learning, independent of initial performance or performance ceiling. The decrease in movement times has been used to quantify learning for several decades (Snoddy, 1926; Crossman, 1959). Several functional forms have been suggested for the fit of movement times (Newell and Rosenbloom, 1981; Heathcote et al., 2000), and variants of an exponential are viewed as the most statistically robust choices (Heathcote et al., 2000). Given the vastly superior number of practiced trials in EXT sequences (Fig. S7), we estimate the *learning rate* for each participant as the average *κ* between both EXT sequences, consistent with previous work (Bassett et al., 2015).

In addition to movement time, we defined *error rate* as the number of incorrect button presses during the full execution of each sequence, and *reaction time* as the time between the onset of a trial and the first button press. We performed a linear fit on both of these additional measures and repeated our main analysis with both their intercept and slope terms (Fig. S4).

### 4.5 MRI data collection

Magnetic resonance images were obtained at 3.0*T* on a Siemens Trio using a 12-channel phased-array head coil. T1-weighted structural images of the whole brain were collected from each subject (repetition time [*TR*] = 15.0 ms; time echo [*TE*] = 4.2 ms; flip angle: 90°; 3D acquisition; field of view: 256 mm, slice thickness: 0.89 mm; 256× 256 acquisition matrix). Data from one resting state run (146 TRs), five experimental runs (variable number of TRs depending on how quickly the task was performed (Bassett et al., 2015)), and a second resting state run (146 TRs) were acquired with a single-shot echo planar imaging sequence that was sensitive to BOLD contrast ([TR] = 2, 000 ms; time echo [TE] = 30 ms; flip angle: 90°; field of view: 192 mm, slice thickness: 3 mm with 0.5 mm gap; 64 × 64 acquisition matrix across 37 axial slices per TR). The present study examines data from four resting state scans, each lasting 5 minutes (150 TRs), acquired at the beginning of each scanning session.

### 4.6 MRI data preprocessing

Cortical reconstruction and volumetric segmentation of the structural data was performed with the Freesurfer image analysis suite (Dale et al., 1999). Preprocessing of the resting state fMRI data involved multiple steps: the first four volumes in each run were discarded to allow stabilization of longitudinal magnetization; sinc-interpolation in time was performed with AFNI’s (Cox, 1996) 3dTshift to correct for the slice acquisition order; orientation of all images was changed to Right-Posterior-Inferior using AFNI’s 3dresample; images were rigid-body motion corrected with AFNI’s 3dvolreg by aligning all volumes with the mean volume (estimated with AFNI’s 3dTstat) in each run; coregistration between the structural image and the mean functional image was performed with Freesurfer’s bbregister (Greve and Fischl, 2009); brain-extracted functional images were obtained by applying Freesurfer’s brain mask on to images from each functional run using AFNI’s 3dcalc; global intensity normalization was performed across all functional volumes using FSL’s fslmaths (Smith et al., 2004) to ensure that all time series were in the same units; functional data was smoothed in surface space with an isotropic Gaussian kernel of 5-mm full width at half-maximum and in the volumetric space with an isotropic Gaussian kernel of 5-mm full width at half-maximum and, using Freesurfer’s `mris_volsmooth`; six motion parameters estimated using Artifact Detection Tools (ART) (Whitfield-Gabrieli, 2009) — three for translation and three for rotation —, as well as the temporal derivatives, quadratic terms, and temporal derivatives of the quadratic terms had their contribution removed from the BOLD signal; non-neuronal sources of noise (white-matter and CSF signals) were estimated by averaging signals within masks obtained with Freesurfer segmentation tools and by identifying voxel time series with high temporal standard deviations, and removed using the anatomical (aCompCor) and temporal CompCor (tCompCor) methods (Behzadi et al., 2007); finally, a temporal band-pass filter of 0.01 Hz to 0.1 Hz was applied using AFNI’s 3dFourier. Motion-censoring was not performed to ensure an equal amount of data per subject.

Using the above processing pipeline, we expect to have been able to correct for motion effects due to volume-to-volume fluctuations relative to the first volume in a scan run. After this motion correction procedure, we observed no correlation between any of the six motion parameters (x-translation, y-translation, z-translation, roll, pitch, and yaw, calculated for each run and training session) and visual-motor connectivity (*P* > 0.05) across all scanning sessions. These results indicated that individual differences in motion were unlikely to drive the effects reported here.

### 4.7 Parcellation scheme

We used a functionally-derived parcellation scheme with 333 regions of interest (ROIs) representing putative cortical areas (Gordon et al., 2014). Region boundaries were identified such that each parcel had a highly homogeneous resting-state functional connectivity pattern, indicating that they contained one unique RSFC signal. We also used a surface-based parcellation of human cortical gyri and sulci (`aparc.a2009s.annot` in Freesurfer) (Destrieux et al., 2010).

In a supplementary set of analyses, we used a volumetric-based parcellation scheme composed of 626 ROIs (AAL-626) that was formed by the combination of two separate atlases: (i) an AAL-derived 600-region atlas (Hermundstad et al., 2013, 2014), which subdivides the 90 AAL anatomical regions into regions of roughly similar size via a spatial bisection method, and (ii) a high-resolution probabilistic 26-region atlas of the cerebellum in the anatomical space defined by the MNI152 template, obtained from T1-weighted MRI scans (1-mm isotropic resolution) of 20 healthy young participants (Smith et al., 2004; Woolrich et al., 2009). The combination of these two atlases provided a high-resolution, 626-region atlas of cortical, subcortical, and cerebellar regions. This volumetric atlas, which we call AAL-626 atlas, has been used previously (Bassett et al., 2015).

### 4.8 Functional connectivity estimation

In previous work, analyses of the task data from the same experiment yielded two sets of ROIs based on the high probability that its regions were assigned to the same functional community by time-resolved clustering methods (Bassett et al., 2015). These two sets of regions broadly corresponded to (i) early visual cortex (which has been referred to as the *visual module;* Fig. 2a) and (ii) primary and secondary somato-motor regions (which has been referred to as the *somato-motor module;* Fig. 2a). A list of region labels associated with the two modules is displayed in Table 1. Because this prior work used regions defined anatomically (AAL-626), we first determined the equivalence with the Gordon333 atlas. Specifically, for each region of the AAL-626 atlas, we determined the region of Gordon333 with the largest spatial overlap in MNI152 space. This procedure, in turn, allowed us to identify the visual and somato-motor modules in the Gordon333 atlas. We then extracted the average resting state time series across regions from each of the functional modules, calculated their Spearman’s rank correlation coefficient (a nonparametric measure of statistical dependence between two variables), and applied a Fisher *r*-to-*z* transformation. We refer to this *z*-value as the *visual-motor connectivity*.

Importantly, the removal of various signal components present throughout most of the brain (in particular by the tCompCor method) leads to a shift in the distribution of functional connectivity values, giving rise to negative correlations. We note that, while these approaches substantially improve the robustness of our results by eliminating physiological noise from the data (Lund and Hanson, 2001), our results remain significant with a less stringent noise removal pipeline that does not shift the range of correlation values (Fig. S9).

We confirmed that the modules identified from the task data were also modules at baseline by comparing the modularity quality (Newman, 2006a) of the actual partition with the modularity quality of 10,000 permuted partitions. The modularity quality is given by equation (2):

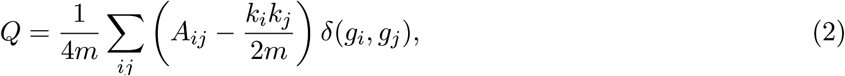

where *A*_*ij*_ is the functional connectivity matrix including all visual and motor regions, *k*_*i*_ and *k*_*j*_ are the strength of nodes *i* and *j, m* = 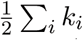 the total strength in the network, and *δ*(*g*_*i*_,*g*_*j*_) = 1 if nodes *i* and *j* belong to the same module or *δ*(*g*_*i*_,*g*_*j*_) = 0 otherwise. We observed that the modularity quality of the actual partition into visual and motor modules was higher than the modularity quality of all 10,000 permuted partitions (*p* = 0.0001), demonstrating that the separation of brain regions into motor and visual modules is an accurate representation of the network organization.

A similar approach was performed for the surface-based analysis, which aimed to identify which specific functional connections within visual and somato-motor areas were most correlated with learning rate. We used broadly defined visual and somato-motor regions of interest (ROIs) and examined the correlations between each visual-to-motor connection and learning rate. The visual ROI was defined as composed of the entire occipital lobe, parieto-occipital, and occipito-temporal areas (Fig. 3a), and the somato-motor ROI was defined as composed of precentral, paracentral, and postcentral sulci and gyri, and central sulcus (Fig. 3a). After projecting the BOLD time-series from each voxel into surface vertices in subject native space, we extracted the average activity within each of the surface-based parcels and calculated the Fisher *r*-to-*z* transform of the Spearman’s rank correlation coefficient between the activity in each region of the visual ROI and each region of the somato-motor ROI.

### 4.9 Measure of statistical relationship

Spearman’s rank correlation was chosen as a measure of statistical relationship between any two variables with different units. This nonparametric statistic measures the extent to which two variables are monotonically related without a requirement for linearity. To assess the relationship between two variables with the same units, Pearson product-moment correlation was used.

## Acknowledgements

DSB acknowledges support from the John D. and Catherine T. MacArthur Foundation, the Alfred P. Sloan Foundation, the Army Research Laboratory through contract no. W911NF-10- 2-0022 from the U.S. Army Research Office, the Army Research Office through contract no. W911NF-14-1-0679, the Office of Naval Research Young Investigator Program, the NIH through award R01-HD086888 and 1R01HD086888-01, and the National Science Foundation awards #BCS-1441502, #BCS-1430087, and #PHY-1554488. The content is solely the responsibility of the authors and does not necessarily represent the official views of any of the funding agencies.

## Author Contributions

NW and SG developed the experimental paradigm. NW collected the data. DB, MM designed the research. MM analyzed the data. MM, NW, AB, GA, and DB wrote the paper. The authors declare no potential conflict of interest.

## Supporting Information

**Figure S1:**
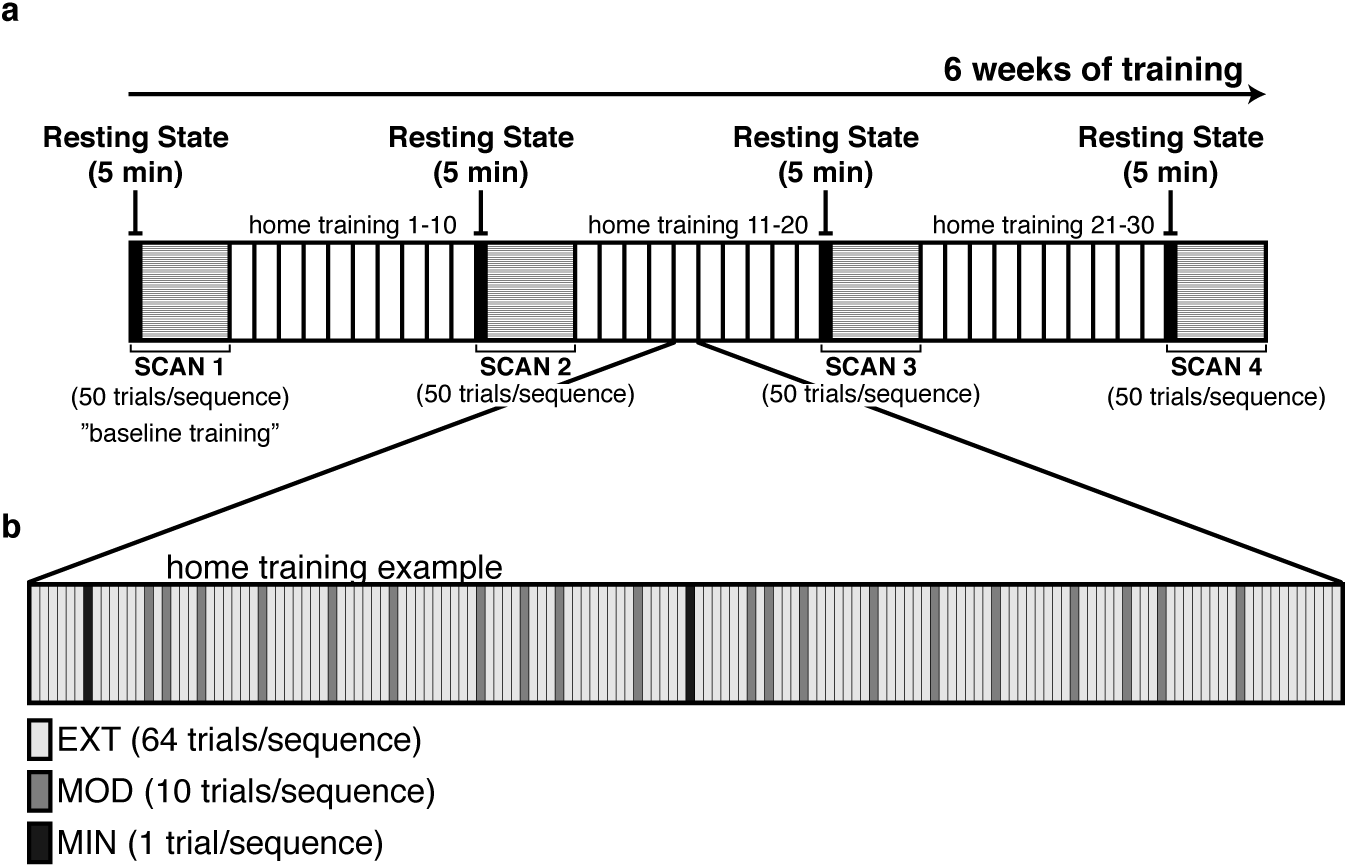
Experiment protocol and timeline. (a) The experiment protocol comprised of six weeks of training of six distinct motor sequences. Following a brief explanation of the task instructions, an initial MRI scan session was held during which blood-oxygen-level dependent (BOLD) signals were acquired from each participant. The scan session began with a resting state scan lasting 5 minutes where participants were instructed to remain awake and with eyes open without fixation. During the remainder of the first scan session (baseline training), participants practiced each of six distinct motor sequences for 50 trials each, or approximately 1.5 hours. They were then instructed to continue practicing the motor sequences at home using a training module that was installed by the experimenter (N.F.W.) on their personal laptops. Participants completed a minimum of 30 home training sessions, which were interleaved with two additional scan sessions, each occurring after at least 10 home training sessions. A final scan session was held following the completion of the 6 weeks of training. The same protocol was followed in each of the four scan sessions: a 5 minute resting state scan, followed by approximately 1.5 hours of the DSP task, where each of six distinct motor sequences was practiced for 50 trials each. (b) Most of the motor sequence training occurred at home, between scanning sessions. An ideal home training session consisted of 150 trials with sequences practiced in random order (randomization used the Mersenne Twister algorithm of Nishimura and Matsumoto as implemented in the random-number generator rand.m of MATLAB version 7.1). Each EXT sequence was practiced for 64 trials, each MOD sequence was practiced for 10 trials, and each MIN sequence was practiced for 1 trial.

**Figure S2:**
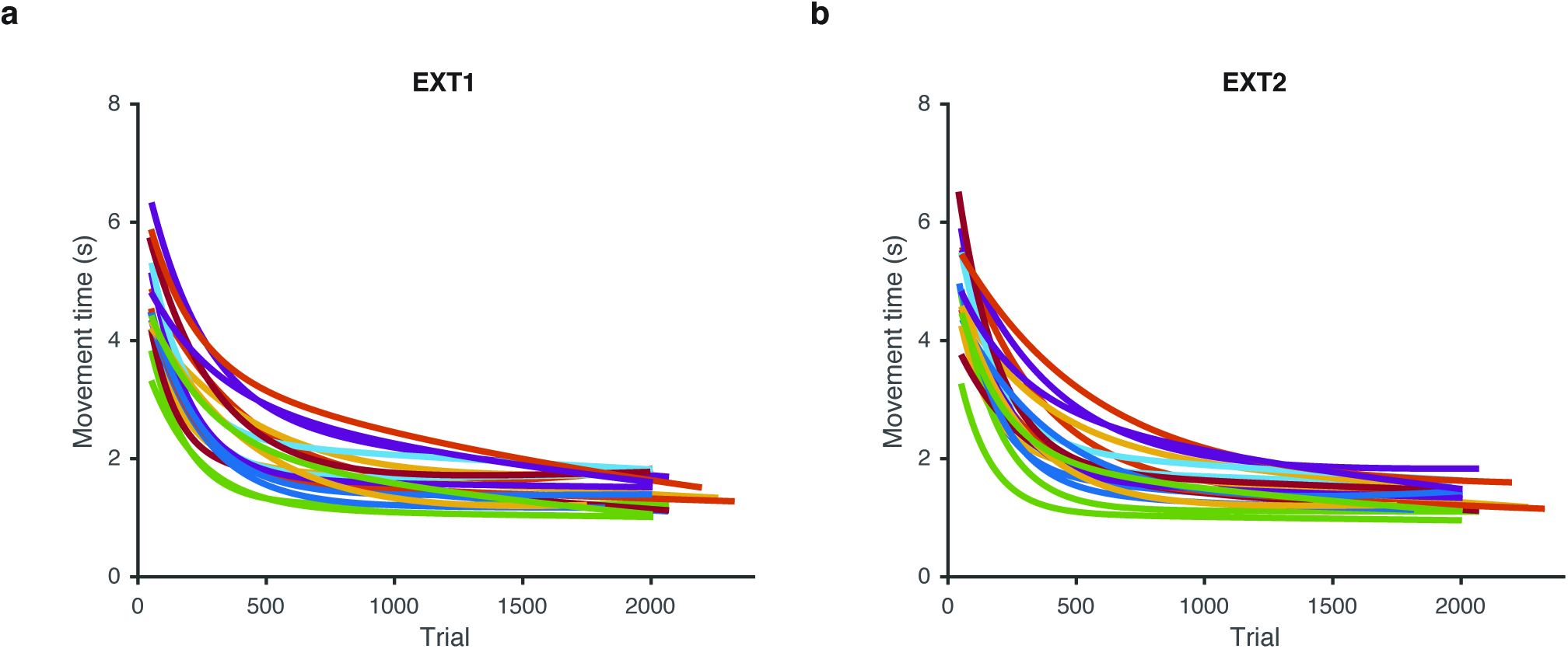
Learning curves from individual participants. Time required to execute a complete motor sequence (*movement time*), as a function of trial number. Colored curves are two-term exponential fits of the movement times from each participant. Learning happened for all participants, as evidenced by the reduction of movement times, but with large variability in the decay rates. (a) EXT1 sequence. (b) EXT2 sequence.

**Figure S3:**
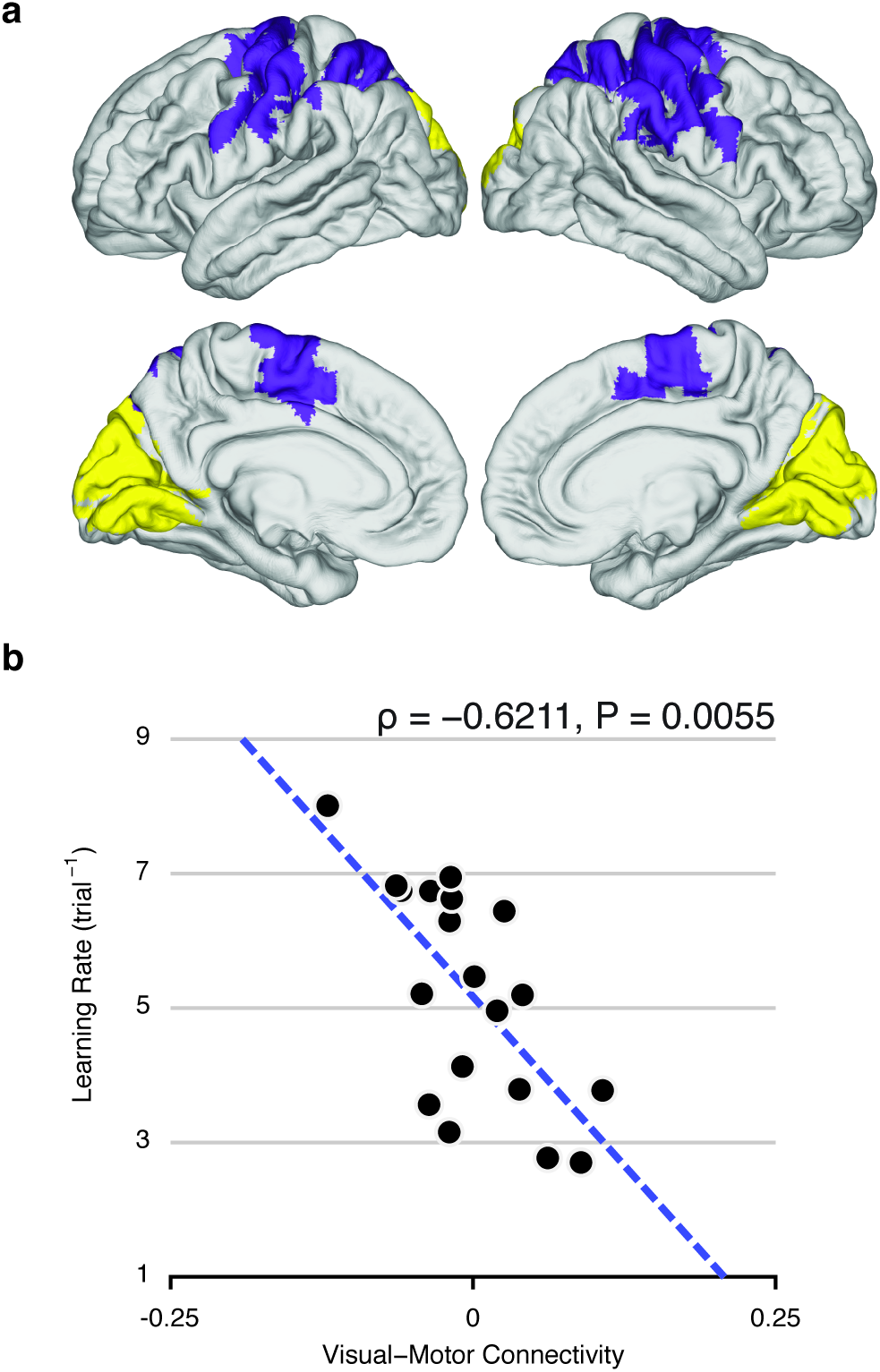
Replication of Figure 2 using an anatomical parcellation (AAL-626). (*a*) Visual module (yellow) and somato-motor module (purple), identified by time-resolved clustering methods applied to BOLD activity acquired during the execution of motor sequences (Bassett et al., 2015). The modules were defined in a data-driven manner and correspond broadly but not exactly to putative visual and somato-motor modules. (*b*) Functional connectivity between visual and somato-motor modules, estimated at rest and prior to learning, reliably predicts individual differences in future learning rate. We define the learning rate as the exponential drop-off parameter of the participant’s movement time as a function of trials practiced, and we define functional connectivity as the average correlation value between activity in visual regions and somato-motor regions.

**Figure S4:**
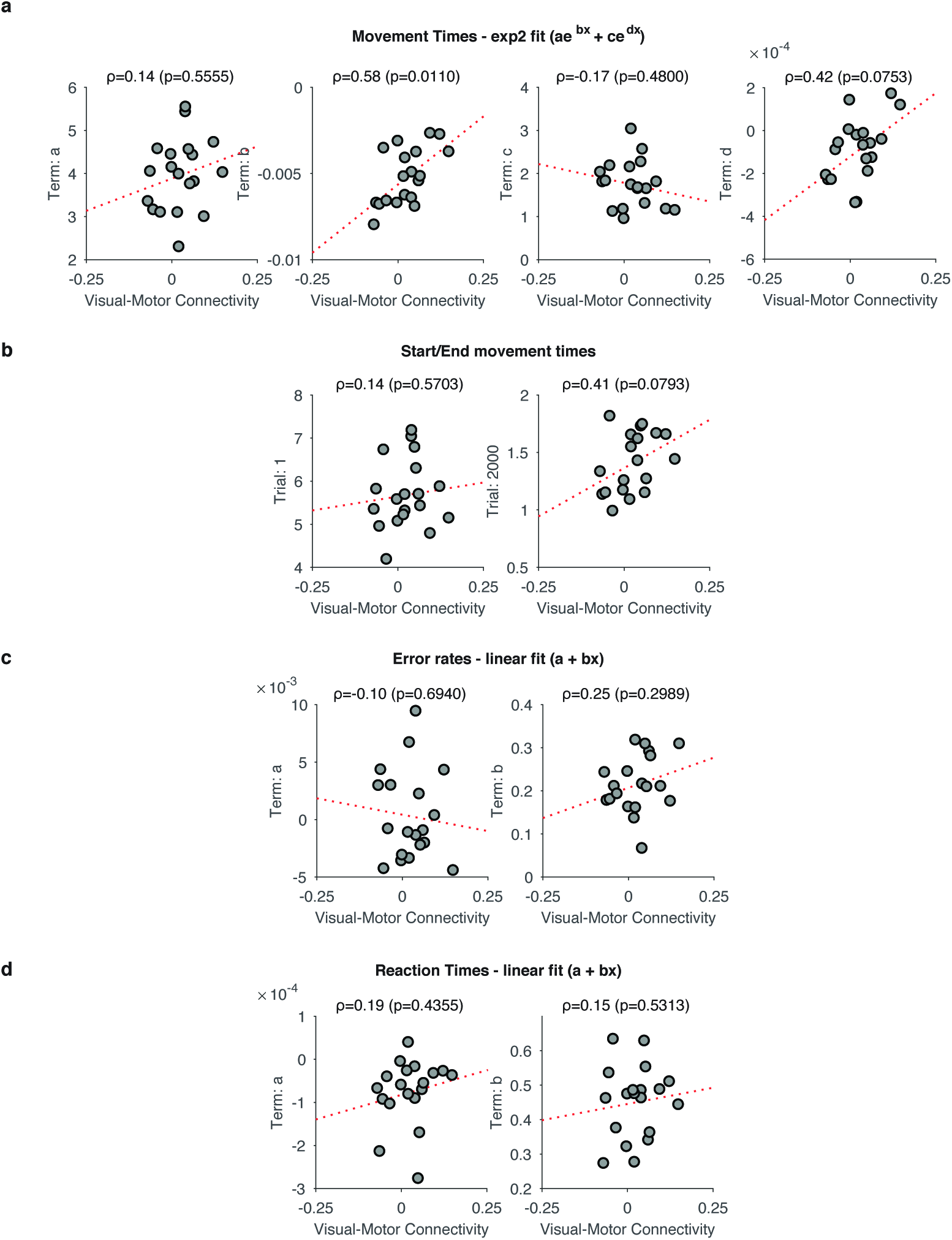
Statistical relationship between resting visual-motor connectivity and different behavioral markers. (a) Relationship between resting-state visual-motor connectivity estimated from the resting-state scan acquired in SESSION 1 and each of the four parameters from the two-term exponential fits of the movement times. Notice the marginal significance of the correlation between visual-motor connectivity and *term d*, suggesting that visual-motor connectivity correlates not only with the faster drop-off parameter (*term b*), but also with the slower decay parameter (*term d*). (b) Relationship between resting-state visual-motor connectivity estimated from the resting-state scan acquired in SESSION 1 and the fitted start movement time (*left*); similarly for fitted end movement time (*right*). Notice the marginal significance of the correlation between visual-motor connectivity and movement time at trial 2000, suggesting that participants with high visual-motor connectivity tend to have longer movement times at the end of the training session. (c) Relationship between resting-state visual-motor connectivity estimated from the resting-state scan acquired in SESSION 1 and both parameters from a linear fit to the error rates. (d) Relationship between resting-state visual-motor connectivity estimated from the resting-state scan acquired in SESSION 1 and both parameters from a linear fit to the reaction times.

**Figure S5:**
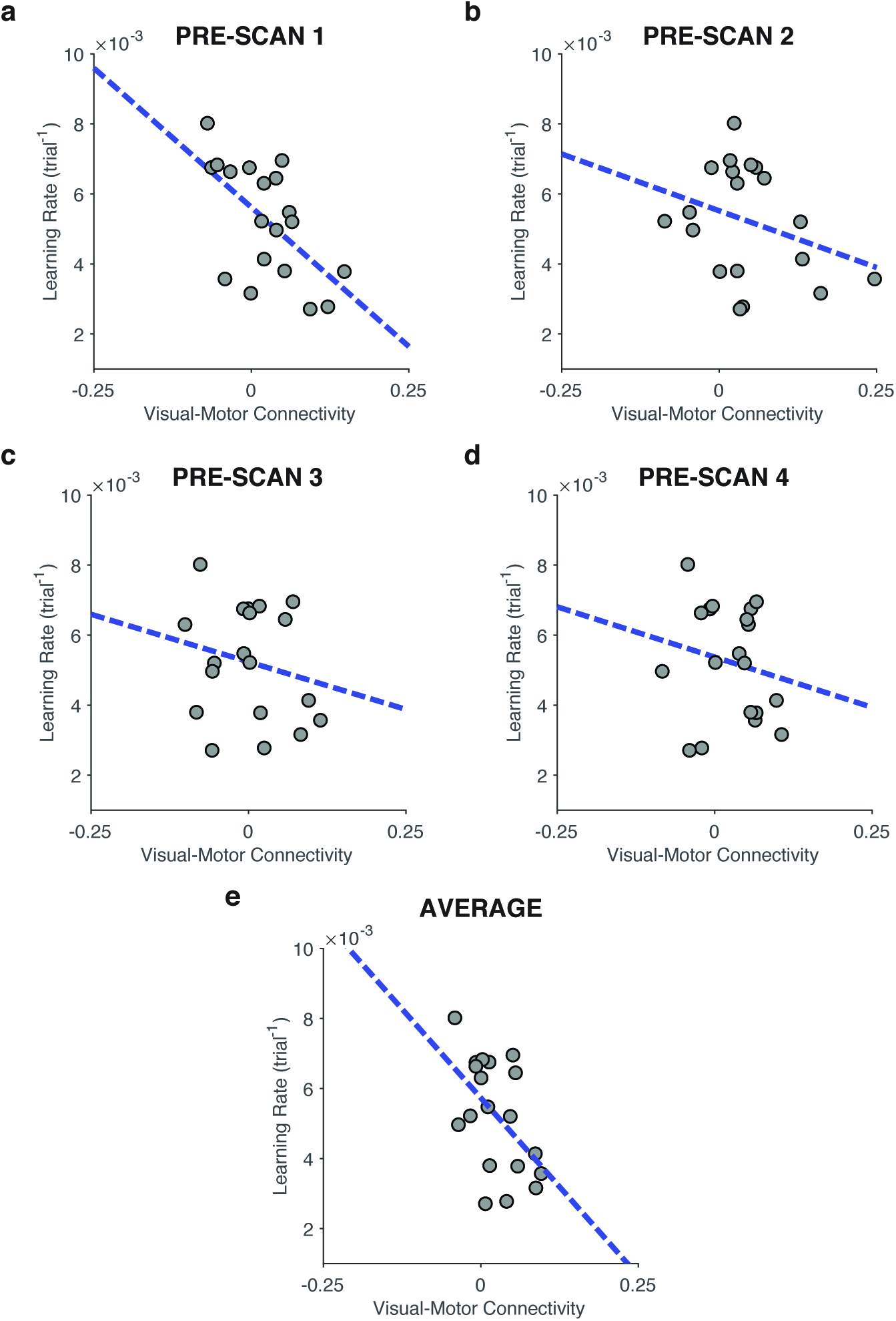
Correlation between visual-motor connectivity at various sessions and overall learning rate. (a) Relationship between visual-motor connectivity estimated from the resting-state scan acquired in SESSION 1 and overall learning rate. The Spearman correlation between these two quantities is *ρ* = −0.5772, *P* = 0.0110. (b) Relationship between visual-motor connectivity estimated from the resting-state scan acquired in SESSION 2 and overall learning rate. The Spearman correlation between these two quantities is *ρ* = −0.2895, *P* = 0.2286. (c) Relationship between visual-motor connectivity estimated from the resting-state scan acquired in SESSION 3 and overall learning rate. The Spearman correlation between these two quantities is *ρ* = −0.1772, *P* = 0.4664. (d) Relationship between visual-motor connectivity estimated from the resting-state scan acquired in SESSION 4 and overall learning rate. The Spearman correlation between these two quantities is *ρ* = −0.1561, *P* = 0.5218. (e) Relationship between average visual-motor connectivity across all four sessions and overall learning rate.The Spearman correlation between these two quantities is *ρ* = −0.4614, *P* = 0.0484.

**Figure S6:**
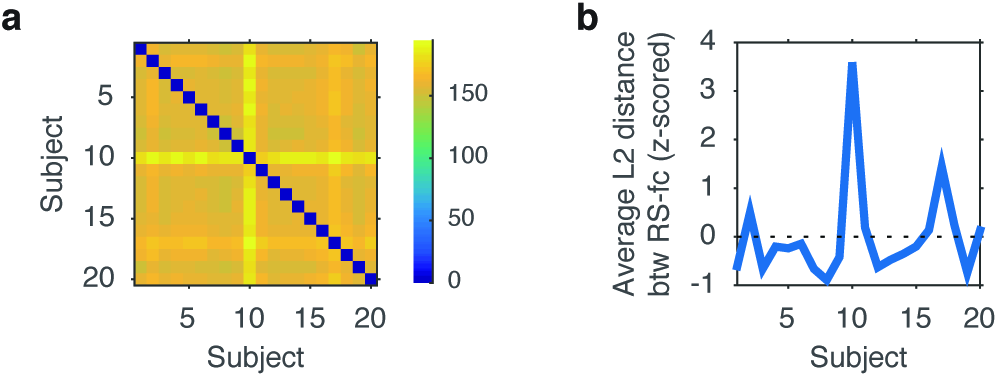
Subject exclusion criterion. (a) We examined resting-state data quality by tracking functional connectivity outliers from our group norm. We calculated the average L2 distance between corresponding cells of the 626 × 626 functional connectivity matrices from all pairs of participants, summarized in the dissimilarity matrix of the figure. (b) Average L2 distance between the RSFC matrix of one participant and that from all others. With the exception of subject 10, all subjects were within 1.5 standard deviations from each other. The resting state data from subject 10 differed on average by 3.6 standard deviations from the others and, therefore, was excluded from the remainder of the analyses.

**Figure S7:**
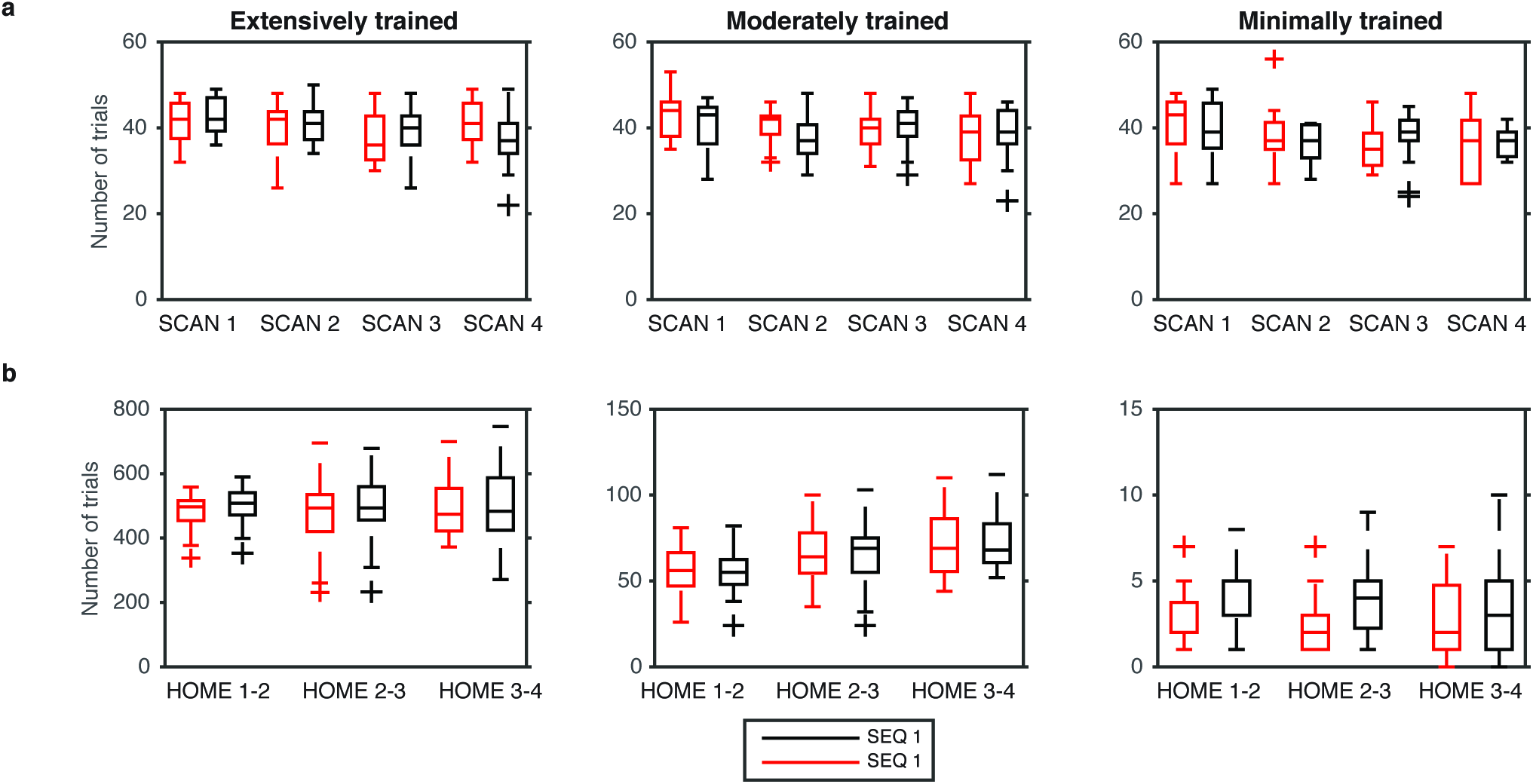
Number of error-free trials performed per session. (a) Number of trials practiced in each scan session. *Left panel*: Extensive training (EXT) session; *Middle panel*: Moderate training (MOD) session; *Right panel*: Minimal training (MIN) session. Box plot represents quartiles and the ‘+’ symbols represent outliers. The variability in the number of executed trials during scan sessions arose mainly due to software or hardware difficulties. (b) Number of trials practiced in each home session. *Left panel*: Extensive training (EXT) session; *Middle panel*: Moderate training (MOD) session; *Right panel*: Minimal training (MIN) session. Box plot represents quartiles and the ‘+’ symbols represent outliers. The variability in the number of executed trials is due to some subjects training more days than others between successive scanning sessions.

**Figure S8:**
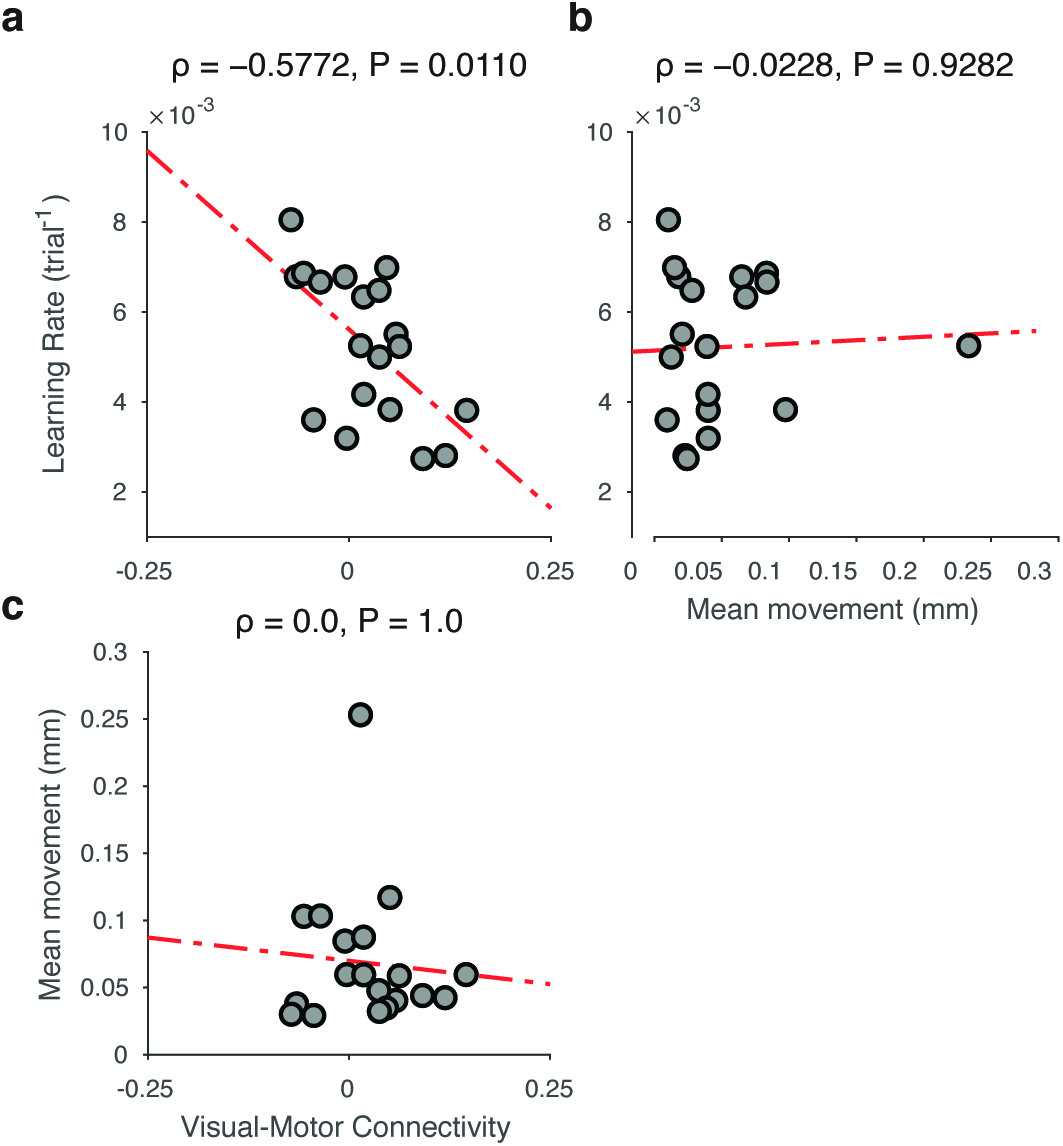
Pairwise relationships between mean subject motion, visual-motor connectivity, and learning rate. (a) Relationship between visual-motor connectivity (in session 1) and learning rate — as in Fig. 2. (b) Relationship between mean subject motion (in session 1) and learning rate. Learning rate was unrelated to subject motion. (c) Relationship between visual-motor connectivity (in session 1) and mean subject motion. Visual-motor connectivity was unrelated to subject motion.

**Figure S9:**
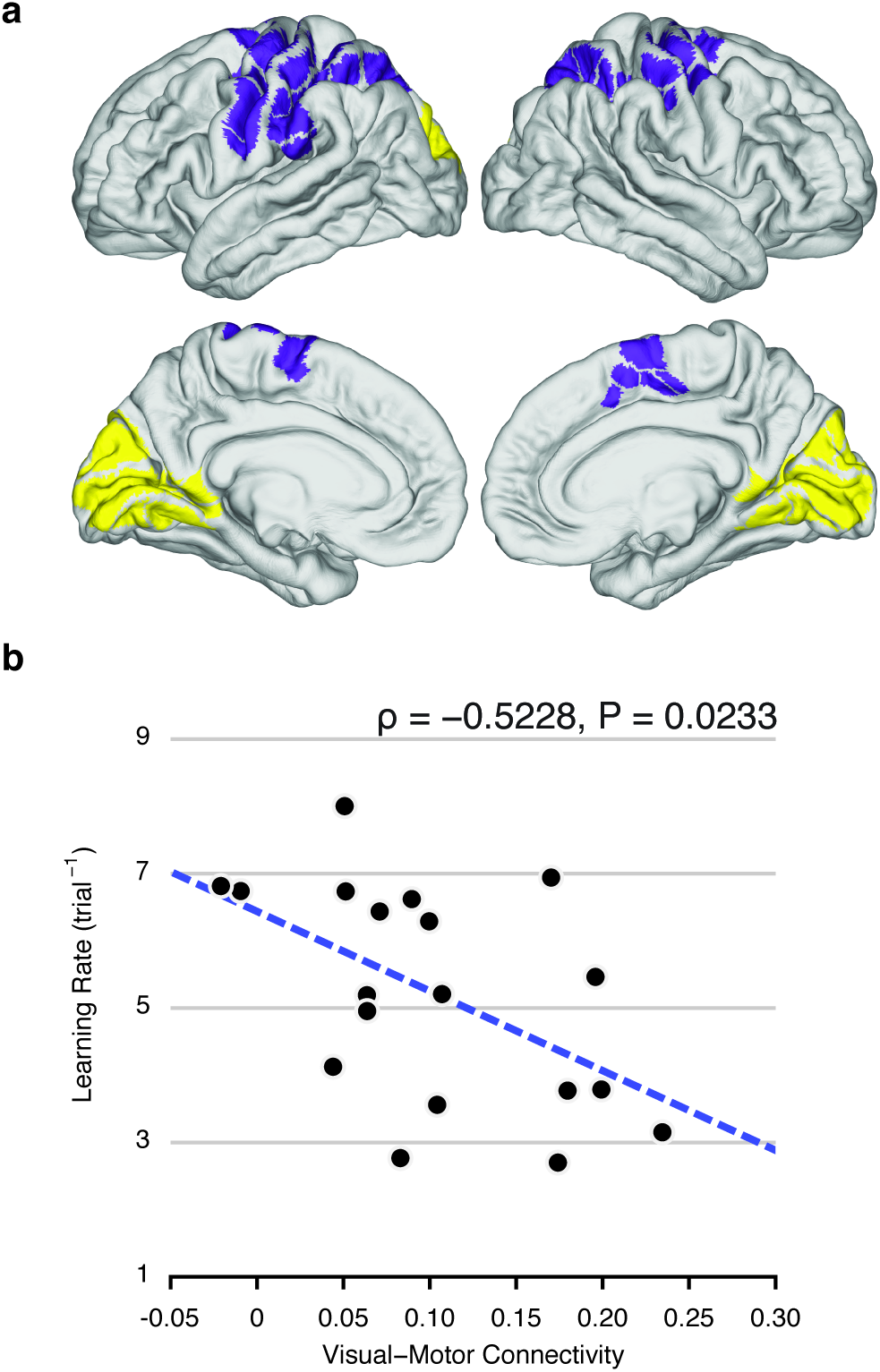
Replication of Fig. 2 with uncentered functional connectivity values. (a) Same as Fig. 2a: Visual (yellow) and somato-motor (purple) modules. (b) Similar to Fig. 2b. The removal of various signal components present throughout most of the brain (in particular by the *tCompCor* method) leads to a shift of the distribution of functional connectivity values, giving rise to negative correlations (Fig. 2b). Here, we use a less stringent noise removal pipeline (same as the original but without the *tCompCor* method) that produces a smaller shift of the range of correlation values. In line with our original results, we observe that functional connectivity between visual and somato-motor modules, estimated at rest and prior to learning, reliably predicts individual differences in future learning rate (*ρ* = −0.5280, *P* = 0.02174). The slightly weaker statistical relationship is likely a consequence of residual physiological noise (Lund and Hanson, 2001).

**Figure S10:**
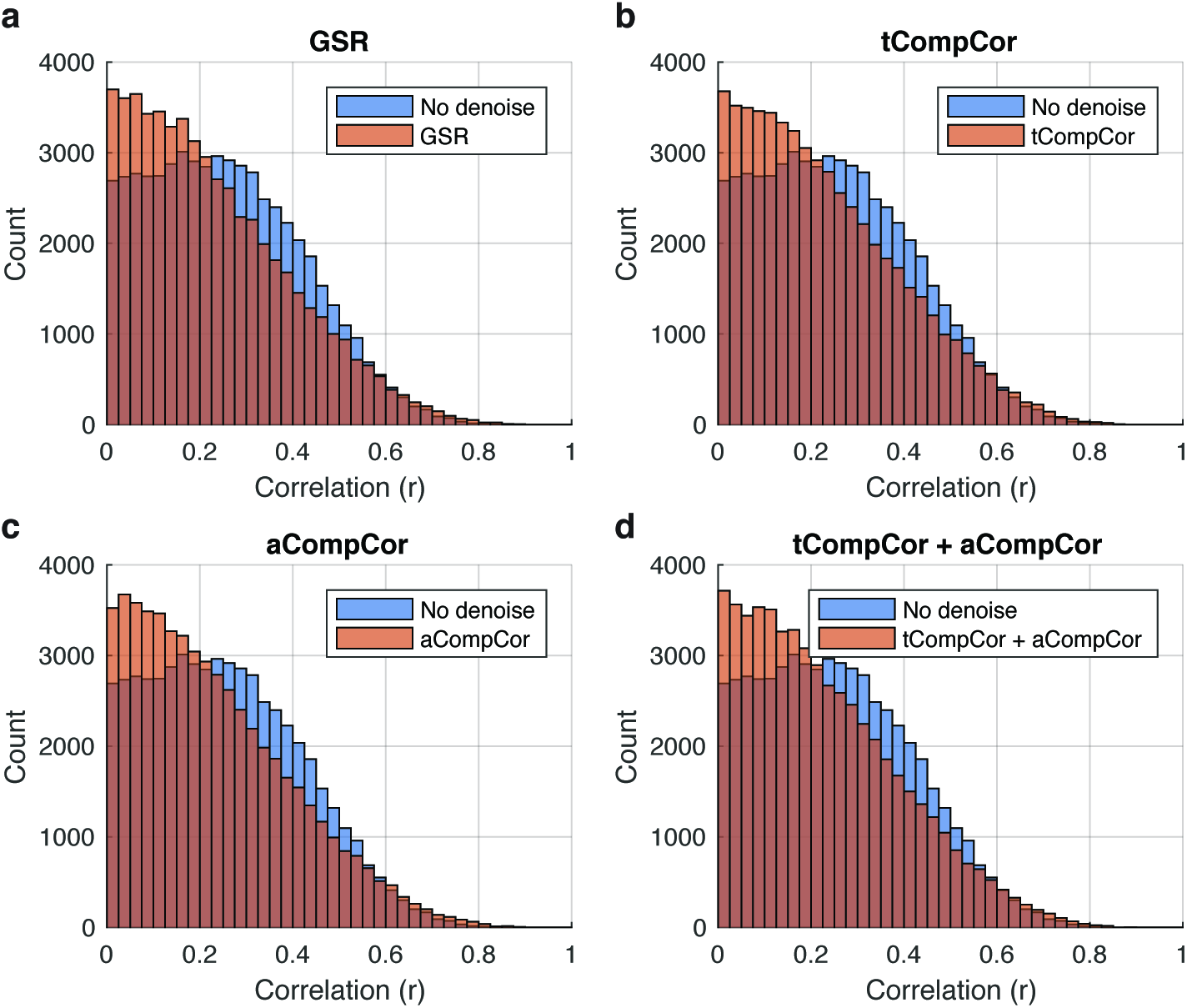
Distribution of correlation values between individual differences in subject mean motion and edge weight for different preprocessing procedures. (a) Global-signal regression (GSR). (b) tCompCor (Behzadi et al., 2007). (c) aCompCor (Behzadi et al., 2007). (d) A combination of tCompCor and aCompCor (Behzadi et al., 2007).

